# HOTAIR promotes an epithelial-to-mesenchymal transition through relocation of the histone demethylase Lsd1

**DOI:** 10.1101/724948

**Authors:** Julien Jarroux, Claire Bertrand, Marc Gabriel, Dominika Foretek, Zohra Saci, Arturo Londoño-Valejo, Marina Pinskaya, Antonin Morillon

## Abstract

Epithelial-to-mesenchymal transition (EMT) drives a loss of epithelial traits by neoplastic cells enabling metastasis and recurrence in cancer. HOTAIR emerged as one of the most renowned long noncoding RNAs promoting EMT mostly as a scaffold for PRC2 and repressive histone H3 Lys27 methylation at gene promoters. In addition to PRC2, HOTAIR interacts with the Lsd1 lysine demethylase, a known epigenetic regulator of cell fate during development and differentiation. However, Lsd1 role in HOTAIR function is still poorly understood. Here, through expression of truncated variants of HOTAIR, we revealed that, in contrast to PRC2, its Lsd1-interacting domain is essential for acquisition of migratory properties by epithelial cells. HOTAIR induces Lsd1 relocation from its inherent genomic loci hence reprogramming the epithelial transcriptome. Our results uncovered an unexpected role of HOTAIR in EMT as an Lsd1 effector and pointed to the importance of Lsd1 as a guardian of the epithelial identity.

## INTRODUCTION

The epithelial-to-mesenchymal transition (EMT) allows normal or neoplastic cells to gradually lose their differentiated epithelial characteristics including cell adhesion and polarity and to acquire mesenchymal traits enabling cytoskeleton reorganization and motility (Lamouille et al., 2014). EMT is closely linked to carcinogenesis since it progressively endows epithelial cells with multiple properties required for invasion and metastasis, but also for acquisition of stem-like properties contributing to tumor recurrence and drug resistance (Ye and Weinberg, 2015). This dynamic and reversible process is driven by complex changes in signaling circuits and reprogramming of gene expression. Transcriptional factors such as zinc-finger E-box-binding (ZEB), Slug, Snail and Twist have been identified as master regulators of EMT, coordinating repression of epithelial genes and activation of mesenchymal genes (Lamouille et al., 2014). The expression and action of these transcription factors can, in turn, be regulated at post-transcriptional level by RNAi pathway (Lamouille et al., 2013), but also epigenetically. In the latter case, chromatin-modifying complexes, such as histone lysine methyltransferases (Polycomb), histone deacetylases (NuRD) and demethylases (Lsd1, PHF2) may determine the transcriptional activity of a genomic locus through covalent chromatin modifications and, as a consequence, may govern the epithelial-mesenchymal plasticity (Tam and Weinberg, 2013). In particular, epigenetic landscape and balance in expression of EMT genes may contribute to the residency of cells in an epithelial state, preserving or maintaining epithelial identity or, in the contrast, allowing transition to a mesenchymal state.

An increasing number of examples supports the involvement of long noncoding (lnc)RNAs in the EMT and metastasis (Huarte and Marín-Béjar, 2015), (Liang et al., 2018), (Shi et al., 2015), (Richards et al., 2015). These RNA polymerase II transcripts of at least 200 nucleotides long and of any or low coding potential can intervene in regulation of gene expression in the nucleus through RNA-protein or RNA-DNA pairing mechanisms, scaffolding and guiding chromatin modifying complexes to specific genomic locations (Quinn and Chang, 2015), (Hendrickson et al., 2016), (Morlando et al., 2014). LncRNAs often show cell- and tissue specific expression and are highly deregulated in cancers. However, molecular mechanisms underlying lncRNAs dysregulation and action remain largely unknown. Among the most prevalent cancer-associated lncRNAs is HOTAIR (for HOX transcript antisense intergenic RNA). Clinical studies have clearly shown its overexpression in most human cancers and its association with poor prognosis, metastasis and acquisition of stemness (Tsai et al., 2011), (Kogo et al., 2011a), (Li et al., 2017), (Gupta et al., 2010), (Zhang et al., 2015). HOTAIR has been firstly identified in human fibroblasts as a molecular scaffold RNA, responsible for epigenetic regulation of cell fate during differentiation (Rinn et al., 2007). Indeed, the majority of nuclear HOTAIR functions have been attributed to its interaction with the Polycomb repressive complex 2 (PRC2) and Histone H3 Lys27 (H3K27) methylation of EMT genes promoters in *trans* (Kogo et al., 2011), (Gupta et al., 2010). If the exact mode of the lncRNA targeting to genomic loci still stays unclear, the molecular outputs are highly cell-type specific. In hepatocytes, HOTAIR has been reported to mediate a physical interaction between the Snail1 transcription repressor and the Enhancer of Zeste Homolog 2 (EZH2) subunit of PRC2, guiding both to specific loci for regulation of hepatocyte *trans*-differentiation program (Battistelli et al., 2016). However, some publications have also demonstrated that PRC2 promiscuously interacts with many structured coding and noncoding RNAs and have claimed PRC2 dispensability for HOTAIR-mediated transcriptional repression (Kaneko et al., 2013), (Kaneko et al., 2014), (Portoso et al., 2017). Instead, HOTAIR have been proposed to play a role in anchoring PRC2 at specific repressed loci, though the ultimate action of HOTAIR and its protein cofactors are still not fully depicted.

The full length HOTAIR is 2.1 nucleotides long and has a modular secondary structure (Somarowthu et al., 2015). In addition to PRC2 binding to first 300 nucleotides of the Domain 1, HOTAIR within its last 500 nucleotides contains another independent domain associated with the Lsd1/REST/CoREST complex (Wu et al., 2013), (Tsai et al., 2010), (Somarowthu et al., 2015). Lsd1/KDM1A, the lysine specific demethylase-1, has been proposed to demethylate H3K4me2 and to reinforce HOTAIR/PRC2-mediated repression of transcription. Chromatin immunoprecipitation (ChIP) and isolation by RNA purification (ChIRP) allowed identification of GC-rich regions of Lsd1 binding sites and a GA-rich consensus sequence for HOTAIR targeting in epithelial cancer cells (Tsai et al., 2010), (Chu et al., 2011). However, little is known of whether and how Lsd1 contributes to HOTAIR action. Lsd1 is a well-known epigenetic regulator of EMT and cancer with, in few cases, a tumor suppression function (Wang et al., 2009), but mostly playing an oncogenic role (Hino et al., 2016), (Harris et al., 2012), (Sun et al., 2016), (Lim et al., 2010), (Schenk et al., 2012), (Feng et al., 2016). The functional duality of Lsd1 can be attributed to the versatility of its substrates and of Lsd1 interacting partners in different biological contexts (Shi et al., 2004), (Metzger et al., 2005), (McDonald et al., 2011). Indeed, in mouse hepatocytes Lsd1 was reported to control the establishment of large organized heterochromatin H3K9 and H3K4 domains (LOCKs) across the genome during EMT. Large scale immunoprecipitation has revealed that Lsd1 interacts with REST/coREST co-repressors in differentiated epithelial cells, as though in TGFβ treated cells undergoing EMT Lsd1 is mostly associated with transcriptional co-activators including several catenins (McDonald et al., 2011). In addition, non-histone targets, such as p53 and DNMT1, and non-enzymatic, scaffold roles have been proposed for Lsd1, particularly, in regulation of enhancer activity in mammals (Lan et al., 2007a), (Lan et al., 2007b), (Zeng et al., 2016), (Wissmann et al., 2007), (Wang et al., 2001), (Huang et al., 2007), (Scoumanne and Chen, 2007), (Roth et al., 2016). Pharmacological inhibitors of Lsd1 impairing its catalytic activity, unexpectedly, have been shown to act through disruption of its scaffold function, particularly with SNAG domain containing proteins, such as the transcriptional repressor GFI (Maiques-Diaz et al., 2018). Whatever the molecular basis of Lsd1 action may be, the biological outcome depends on a balance between activated and repressed genes underlying the pivotal role of Lsd1 in the phenotypic plasticity of a cell.

In the present study, we aimed to understand a role for HOTAIR interaction with Lsd1 in the EMT reprogramming. For this purpose, we used gain- and loss of function approaches overexpressing HOTAIR in immortalized primary epithelial cells and disrupting HOTAIR interactions with chromatin modifying complexes by deletion of either the 5’-PRC2 or 3’-Lsd1-interacting domains within the lncRNA. As expected, HOTAIR promoted migration of epithelial cells; however, this required the presence of the Lsd1-interacting domain while the PRC2 one was dispensable. At molecular levels, epithelial cells expressing the HOTAIR variant truncated for the Lsd1-interacting domain expressed more and showed less diffused outer membrane distribution of the tight junction protein ZO-1/TJP1, compared to the full-length and the truncated for PRC2 HOTAIR variant. Genome-wide Lsd1 profiling confirmed that the expression of HOTAIR with the intact 3’-extremity induces dramatic changes in chromatin distribution of Lsd1. We propose that HOTAIR, when expressed in epithelial cells, promotes a displacement of Lsd1 from its inherent targets resulting in transcriptomic changes in favor of mesenchymal traits. Our findings pinpoint Lsd1 as a guardian of epithelial identity and support a PRC2-independent function of HOTAIR in acquisition of migratory properties by epithelial cells at very early steps of carcinogenesis.

## RESULTS

### Generation of Epi cell lines expressing full-length and truncated variants of HOTAIR

To decipher a role of the Lsd1 interacting domain in HOTAIR function, our rational was to generate expression vectors containing the lncRNA as a full-length transcript (HOT), but also truncated for the first 300 or the last 500 nucleotides sequences, previously reported to be involved in PRC2 and Lsd1 interactions (HOTΔP and HOTΔL), respectively (Figure 1A). These constructs were transduced into immortalized human epithelial kidney cells, HA5-Early. This cell line, originally obtained from a primary kidney epithelium by ER-SV40 and hTERT transformation very early in their lifespan, is characterized by normal karyotype and epithelial traits, such as rounded cobblestone morphology, low migration and expression of epithelial markers (zonula occludens-1/ZO-1, β-catenin, claudin-1) (Figure S1A-S1C) (Castro-Vega et al., 2013). To facilitate further reading, the HA5-Early cell line is referred to as Epi, and its derivatives as Epi-CTR, Epi-HOT, Epi-HOTΔP and Epi-HOTΔL.

**Figure 1.**
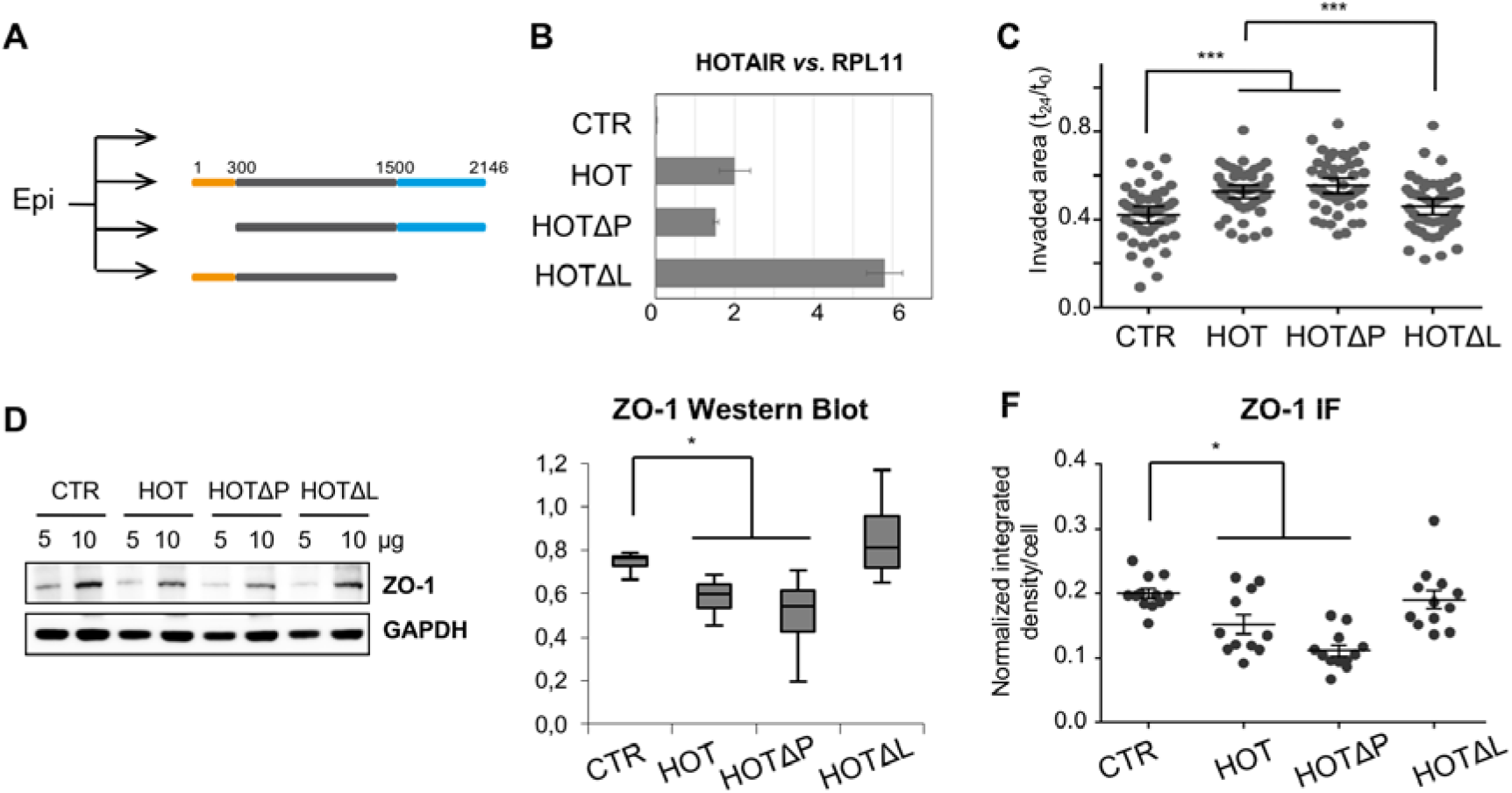
HOTAIR expression in epithelial Epi cells promotes cell migration in Lsd1-dependent manner: **(A)** Stable Epi cell lines overexpressing CTR, full-length and truncated variants of HOTAIR lacking PRC2 or Lsd1-interacting domains, HOTΔP and HOTΔL, respectively; **(B)** Random-primed RT-qPCR measurement of HOTAIR expression in Epi cell lines. cDNA levels are presented as a mean ± standard deviation (SD) for at least three biological replicates; **(C)** Quantification of the wound area invaded in 24 hours by Epi cells as a mean ± Confidence Interval (CI) of 95%; **(D)** Abundance of ZO-1 in Epi cells assessed by Western blot of whole protein extracts; **(E)** ImageJ quantification of ZO-1 in four independent Western blot experiments; **(F)** ZO-1 quantification of IF images performed using the Fiji software and bar-plotted as normalized integrated densities per cell with as a mean ± standard error of the mean (SEM) for at least 11 high-field units representing at least 100 cells; * p-value < 0.05, Student’s t-Test.

Expression levels of all HOTAIR variants were measured relative to the housekeeping protein-coding gene *RPL11* showing stable expression in all experimental conditions. For comparison, we also used the mesenchymal cell line HA5-Late, below referred to as Mes, which derives from the same primary kidney tissue as Epi, but through immortalization at the late steps of the life span after natural accomplishment of EMT (Castro-Vega et al., 2013). In addition to expression of key mesenchymal markers and increased migration properties, this cell line is characterized by high levels of HOTAIR comparing to Epi (Figure S1A-S1C). We found that the ectopic expression of HOTAIR from the CMV promoter was at least 36 times higher comparing to its inherent levels in Epi cells, and 3.5 times higher than in Mes cells expressing it endogenously. While comparing expression levels of the full-length and truncated variants, the HOTΔL transcript was at least twice more abundant than HOT or HOTΔP transcripts in Epi cells (Figure 1B). Subcellular fractionation into cytosolic, nucleoplasm and chromatin fractions confirmed that the overexpression, as well as sequence truncations did not change HOTAIR subcellular residence and, in particular, its association with chromatin in Epi cells in comparison to Mes (Figure S1D, S1E). Moreover, subcellular localization and expression levels of HOTAIR protein partners EZH2 and Lsd1 were the same in all cell lines (Figure S1F and S1G). Generated Epi cell lines expressing the full-length HOT and truncated HOTΔP and HOTΔL were further used as a model system to assess a role of PRC2- and Lsd1-interacting domains in HOTAIR function at cellular and molecular levels.

### Lsd1-interacting domain is essential for HOTAIR function in promoting cell migration

One of the most robust phenotypes associated with HOTAIR expression is the increase in ability of epithelial cells to migrate (Ding et al., 2014), (Dong and Hui, 2016). Therefore, we assessed whether HOTAIR affects migration of Epi cells using the wound healing assay (WHA). As expected, HOTAIR promoted migration of epithelial cells, though the wound healing was much slower than in fully reprogramed mesenchymal Mes cells (Figure 1C, Figure S2A). Surprisingly, deletion of the PRC2-interacting domain did not have an effect as the Epi-HOTΔP cell line migrated as fast as Epi-HOT. On the contrary, HOTAIR missing the Lsd1-interacting domain healed the wound as slowly as the control epithelial cells Epi-CTR (Figure 1C, Figure S2A). We also assessed the proliferation by measuring population doubling (PD) rates of each cell line and did not find significant differences that could explain observed gain or loss of the wound healing efficiency (Table S1).

Cellular migration is a highly complex phenomenon associated with changes in cell-cell junctions, cytoskeletal organization and apico-basal polarity of epithelial cells (Lamouille et al., 2014). We assessed the general morphology of cells expressing HOTAIR by the phalloidin staining of F-actin fibers, but no change in cell shape or in formation of cell sheets was detectable (Figure S2B). During EMT, the acquisition of migratory properties is known to result from the decrease in the formation of tight junctions involved in cell-to-cell contacts (Tornavaca et al., 2015). Therefore, we measured protein levels of the tight junction protein ZO-1/TJP1 by Western blot as well as its subcellular localization by Immunofluorescence (IF). Strikingly, the ZO-1 abundance in the whole protein extracts decreased in HOT and HOTΔP expressing Epi cells showing higher migration, but not in low-migrating Epi-HOTΔL and Epi-CTR cells (Figure 1D and 1E). Concordantly, we observed a more diffused localization of ZO-1 by IF, especially, at cell-cell junctions in HOT and HOTΔP comparing to HOTΔL and CTR (Figure 1F, Figure S2C). The expression of other epithelial markers, β-Catenin and Claudin-1, and mesenchymal markers, Slug, Snail, Zeb1 and Vimentin, was unchanged at protein levels as assessed by Western blot (Figure S3).

Together, these findings suggested that acquisition of migratory properties by epithelial cells is promoted by high levels of HOTAIR and relies on its interaction with Lsd1 rather than with PRC2. The gain in migration is associated amongst other factors with the weakening of cell-cell junctions. The expression of EMT drivers, the key transcription factors known to induce cell reprogramming towards mesenchymal identity, remained unchanged in Epi cells expressing HOTAIR.

### HOTAIR expression in epithelial cells induces global transcriptomic changes, majorly dependent on both PRC2- and Lsd1-interacting domains

To get insights into the molecular mechanisms driving the changes of migratory properties and cell identity upon HOTAIR expression, we performed RNA-sequencing (RNA-seq) and differential expression analysis of CTR, HOT, HOTΔP or HOTΔL expressing Epi cells. First, the transcriptome of Epi-CTR was compared to Epi cells expressing each of HOTAIR variants to define HOTAIR induced transcriptomic changes associated with each condition. Then, differentially expressed (DE) genes of each set were intersected to query the ones common to all, at least to two or exclusive to one specific condition.

First and as expected, HOTAIR expression in Epi cells induced global changes in expression of protein-coding genes (PCGs) with a prevalence of a repressive effect (Figure 2). A total of 743 PCGs were retained as significantly dysregulated in Epi-HOT with a fold-change (FC) above 2 and the adjusted p-value below 0.05 (Figure 2A and 2B). Deletion of either PRC2 or Lsd1-interacting domains within HOTAIR resulted in more moderate transcriptomic perturbations with 191 and 347 DE-PCGs, respectively, again with a prevalence of down-regulation. Further intersection of up- and down-regulated genes associated with each variant identified 495 DE-PCGs genes strictly requiring the presence of both domains (Figure 2B). These genes were grouped into HPL-neg and HPL-pos sets for down- (n=379) and up- (n=116) regulated genes, respectively, representing putative HOTAIR/PRC2/Lsd1-dependent targets (Figure 2B). Some genes of the HPL set have already been reported among EMT markers (Gröger et al., 2012) and some identified as repressed by HOTAIR and PRC2 mediated histone H3 Lysine 27 methylation in previous studies (Gupta et al., 2010). Among them were genes involved in proteolysis of extracellular matrix, *SERPIN2* and *MMP3*, and the protocadherin gene family member *PCDH18* (Gupta et al., 2010), (Xu et al., 2013), (Qiu et al., 2014). Gene ontology (GO) analysis revealed enrichment of the HPL-set for genes involved in several KEGG (Kyoto Encyclopedia of Genes and Genomes) pathways tightly linked to EMT, cancer and metastasis (Figure 2D). Importantly, the most significantly depleted pathways were enriched in genes of extracellular matrix (ECM) receptor interactions, focal adhesion and Hedgehog signaling, whereas the up-regulated genes represented Jak-STAT and bladder cancer pathways (Figure 2D). Notably, the DE-gene sets associated with HOTAIR expression were devoid of the key transcription factors inducing EMT and described as EMT drivers.

**Figure 2.**
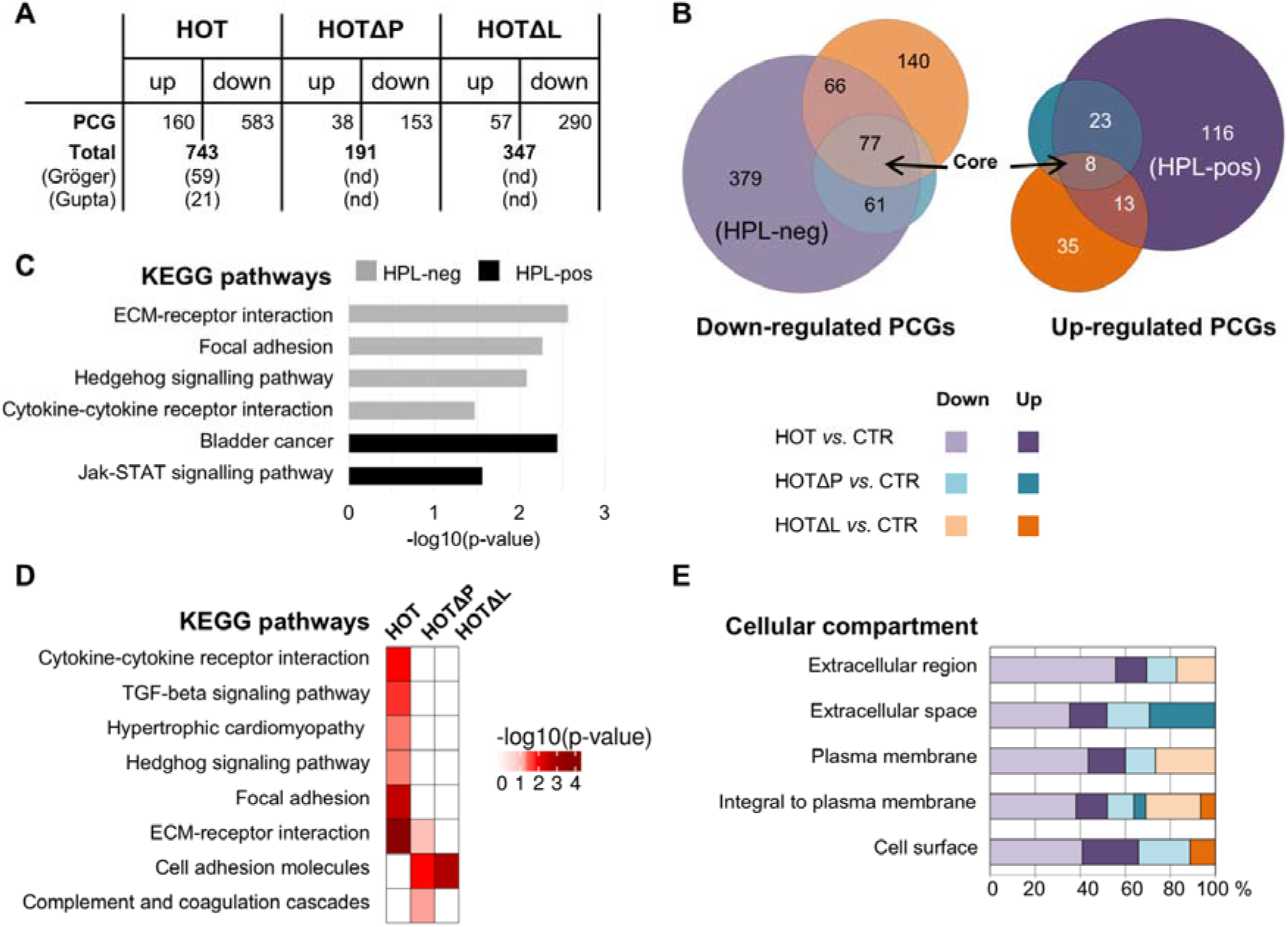
HOTAIR expression in Epi cells induces drastic changes in expression of PCGs, majorly dependent on the presence of both, PRC2- and Lsd1-interacting domains: **(A)** Number of up- and down regulated genes defined as differentially expressed (DE) in HOT, HOTΔP and HOTΔL expressing Epi cells comparing to Epi-CTR by DESeq (FC above 2 and adjusted p-value below 0.05), including those associated with EMT and already identified as HOTAIR/PRC2 targets; nd stands for non-determined; **(B)** Venn diagram of intersection of down- and up-regulated PCGs in Epi-HOT, HOTΔP and HOTΔL cells comparing to Epi-CTR; **(C)** KEGG pathways identified by DAVID as significantly enriched (adj. p-value < 0.05) for HPL-set of PCGs; **(D)** KEGG pathways identified by DAVID as significantly enriched (p-value < 0.05) for DE-PCGs of HOT, HOTΔP and HOTΔL expressing Epi cells comparing to Epi-CTR; **(E)** Cellular compartments of differentially expressed protein counterparts shared by up- and down-regulated PCGs in Epi-HOT, HOTΔP and HOTΔL cells comparing to Epi-CTR.

By contrast, expression of 85 genes was affected whatever the HOTAIR truncation (Figure 2B). These genes were grouped into a Core-set representing potential HOTAIR targets, most likely regulated independently of PRC2 and Lsd1 but through alternative mechanisms. The majority of the Core-set genes was down-regulated (n=77/85) and was involved in protein maturation and processing pathways, proliferation and extracellular matrix organization processes. Since this particular group of genes was not the focus of this study it was excluded from further analysis. Together, our protein and global transcriptomic results strongly suggest that HOTAIR has no function in the EMT reprogramming in immortalized primary epithelial cells, instead, they further reinforce its role as modulator of the epithelial-mesenchymal plasticity towards acquisition of some mesenchymal traits particularly affecting signal transduction and migration pathways.

### Distinct roles of PRC2- and Lsd1-interacting domains in HOTAIR-mediated regulation of gene expression

To further discriminate HOTAIR targets dependent on its interaction with either PRC2 or Lsd1, we analyzed more in detail DE-PCGs in Epi cells expressing truncated variants of HOTAIR. As aforementioned, deletion of either PRC2- or Lsd1-interacting domain within HOTAIR resulted in decreased number of DE genes (R^2^ of 0.986 and 0.982, respectively) comparing to changes induced by the expression of the full-length transcript (R^2^ of 0.972) (Figure 2A, Figure S4).

Paradoxically, even if there were more drastic transcriptome perturbations in Epi-HOTΔL (347 DE-PCGs), they were not sufficient for the Lsd1-domain truncated HOTAIR variant to promote migration, whereas the HOTΔP variant induced less changes (191 DE-PCGs) but still showed increased migration as much as the full-length HOTAIR (Figure 1C). Moreover, while querying GO terms for DE-PCGs in Epi cells expressing truncated variants of HOTAIR, we retrieved cell adhesion molecules for both Epi-HOTΔL and HOTΔP, but for the rest the transcriptomic landscape of these two cell lines was quite distinct (Figure 2D). In addition to KEGG, we searched for cellular compartments of differentially expressed protein counterparts. Strikingly, DE-PCGs of HOT and HOTΔP were particularly enriched in genes localized to cell surface, extracellular region and matrix, as though DE-PCG of HOTΔL were much more represented by plasma membrane and intracellular locations (Figure 2E). Even if GO terms comparisons are difficult to interpret because of the low number of misregulated genes, fast-migrating Epi-HOT and Epi-HOTΔP cells were characterized by overexpression of genes featuring extracellular space and involved in cell adhesion, whereas the low-migrating Epi-CTR and Epi-HOTΔL cells showed more alterations in expression of genes with intracellular functions as cell signaling.

### The LSD1 interacting domain of HOTAIR shapes the fast migrating transcriptome landscape

Epithelial plasticity is defined by a balance in expression of epithelial and mesenchymal genes. Juxtaposition of migration and transcriptome changes in Epi cells overexpressing full-length or truncated variants of HOTAIR strongly suggested that HOTΔL cells maintain their epithelial balance to a larger extent than HOT or HOTΔP cells, both able to interact with Lsd1. This observation nourished a hypothesis that Lsd1/HOTAIR crosstalk may affect epithelial-mesenchymal balance. To define other genes involved in this regulation, we performed a differential expression analysis of Epi-CTR and Epi-HOTΔL transcriptomes against Epi-HOT and Epi-HOTΔP. Knowing that Lsd1 is a component of multiple complexes with repressor or activator activities, we assigned all PCGs that are significantly up-regulated in Epi-CTR and Epi-HOTΔL datasets relatively to Epi-HOT and Epi-HOTΔP (n=148) (DESeq, FC > 1.5 and p-value < 0.05) but absent in the Core-set, as a *Low Migration Signature* (LMS, n=131) (Figure 3A). Similarly, all up-regulated PCGs in Epi-HOT and Epi-HOTΔP (n=77), but absent in the Core-set were grouped into a *High Migration Signature* (HMS, n=75) (Figure 3A).

**Figure 3.**
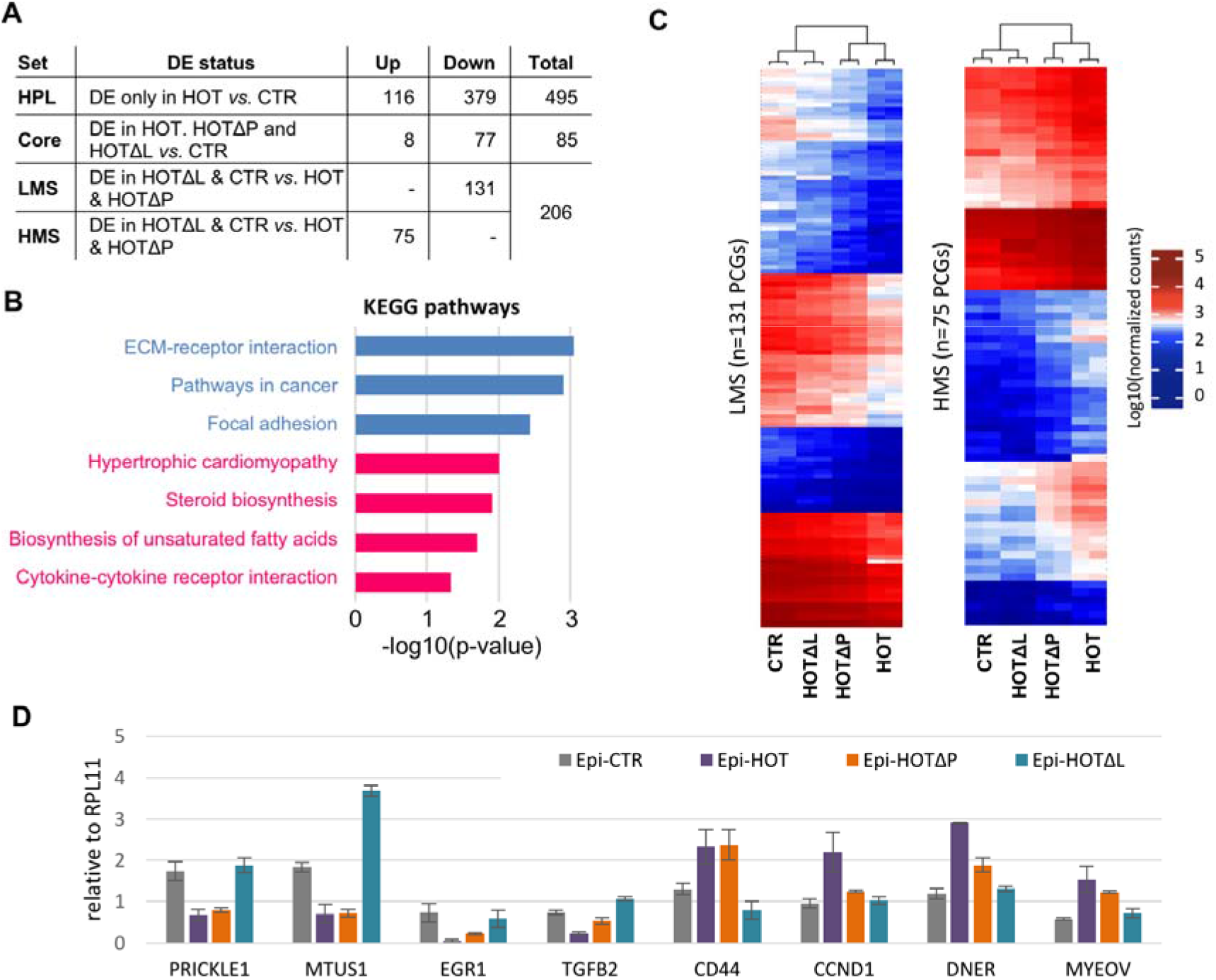
Transcriptome signature of low and fast migrating epithelial cells: **(A)** PCGs assignment to different gene sets according to DE features; **(B)** KEGG pathways enriched by PCGs from LMS and HMS sets (DAVID, p-value below 0.05); **(C)** Unsupervised hierarchical clustering heatmap of LMS and HMS gene sets; **(D)** Random-primed RT-qPCR quantification of gene expression levels relative to RPL11 in Epi-CTR and Epi cells expressing HOT, HOTΔP or HOTΔL variants of HOTAIR.

Remarkably, the LMS-set was composed of genes mostly involved in pathways linked to cardiac development, steroid biosynthesis and cytokine-cytokine receptor interactions, whereas the HMS-set was clearly enriched in cancer and metastasis related pathways including the ECM receptor interaction and focal adhesion (Figure 3B). Notably, transcriptomic changes induced by HOTAIR did not result in a complete switch of EMT program but rather in modulation (attenuation or increase) of gene expression (Figure 3C). Among LMS genes highly expressed in Epi-CTR and Epi-HOTΔL were the tumor suppressor *MTUS1* (Di Benedetto et al., 2006), the nuclear receptor *PRICKLE1/RILP* implicated in the nuclear trafficking of REST/NRSF and REST4 transcription repressors (Shimojo and Hersh, 2006). HMS genes highly expressed in Epi-HOT and Epi-HOTΔP included the cell cycle regulator *CCND1*, the *DNER* activator of the NOTCH pathway, but also the *CD44* EMT marker (Figure 3D).

In conclusion, disruption of HOTAIR interaction with PRC2 or Lsd1 does not abolish completely its function as a regulator of gene expression; however, HOTAIR association with Lsd1 is essential for the modulation of the transcriptomic pattern of epithelial cells in favor of mesenchymal identity.

### The Lsd1-interacting domain of HOTAIR is essential for Lsd1 chromatin redistribution

With the support of previous studies, we hypothesized that the epithelial-mesenchymal plasticity is controlled by the function of Lsd1 in gene expression regulation, which may be modulated in cells expressing HOTAIR. Since Lsd1 protein levels in total and nuclear cell extracts did not show any changes in response to HOTAIR overexpression in epithelial cells (Figure S1F and S1G), we aimed to test whether HOTAIR would affect Lsd1 chromatin occupancy and distribution in two phenotypically distinct groups: Epi-CTR and Epi-HOTΔL cells would represent a biological context, in which Lsd1 exhibits its function independently of HOTAIR maintaining epithelial identity and low migration, whereas Epi-HOT and Epi-HOTΔP would designate a context with both free and HOTAIR associated Lsd1 function. We performed a Chromatin ImmunoPrecipitation sequencing (ChIP-seq) of Lsd1 in Epi cells expressing full-length or truncated variants of HOTAIR in comparison to the control Epi-CTR condition to define Lsd1 chromatin occupancy. The uniquely mapped reads of two replicates per condition were subjected independently to a peak calling procedure of SICER, an algorithm specifically designed for identification of dispersed IP-DNA enriched islands relative to a corresponding Input-DNA signal (Zang et al., 2009). The blacklisted by ENCODE genomic regions were excluded from further consideration (ENCODE Project Consortium, 2012) and only peaks showing at least 1 nucleotide overlap in two replicates were merged and retained for further analysis (Figure 4A). The number of detected Lsd1 peaks was strikingly heterogeneous between conditions, in particularly, CTR and HOTΔL datasets showed as much as 20 times more peaks than HOT and HOTΔP regardless the identical ChIP-seq metrics (Figure S5A, S5B). This result correlated with the differences in migration capacities of the cell lines; CTR and HOTΔL showing lower and HOT and HOTΔP higher migration. We interrogated genomic locations occupied by Lsd1 in both replicates and revealed that only few peaks were located to promoter regions, transcriptional start sites (TSS) as though the majority was detected within noncoding 5’UTR, intergenic and intronic regions (Figure 4A, Figure S5B). Herein, the Epi-HOTΔP cell line was particularly depleted for Lsd1 in promoter and 5’UTR regions. However, no specific, discriminating feature was found when comparing Lsd1 peak locations between the two phenotypically distinct groups, CTR and HOTΔL *versus* HOT and HOTΔP. Distribution of Lsd1 peaks as a distance from genes TSS did not show any significant difference between CTR and three other conditions (p-value > 0.3, Wilcoxon test) (Figure S5C).

**Figure 4.**
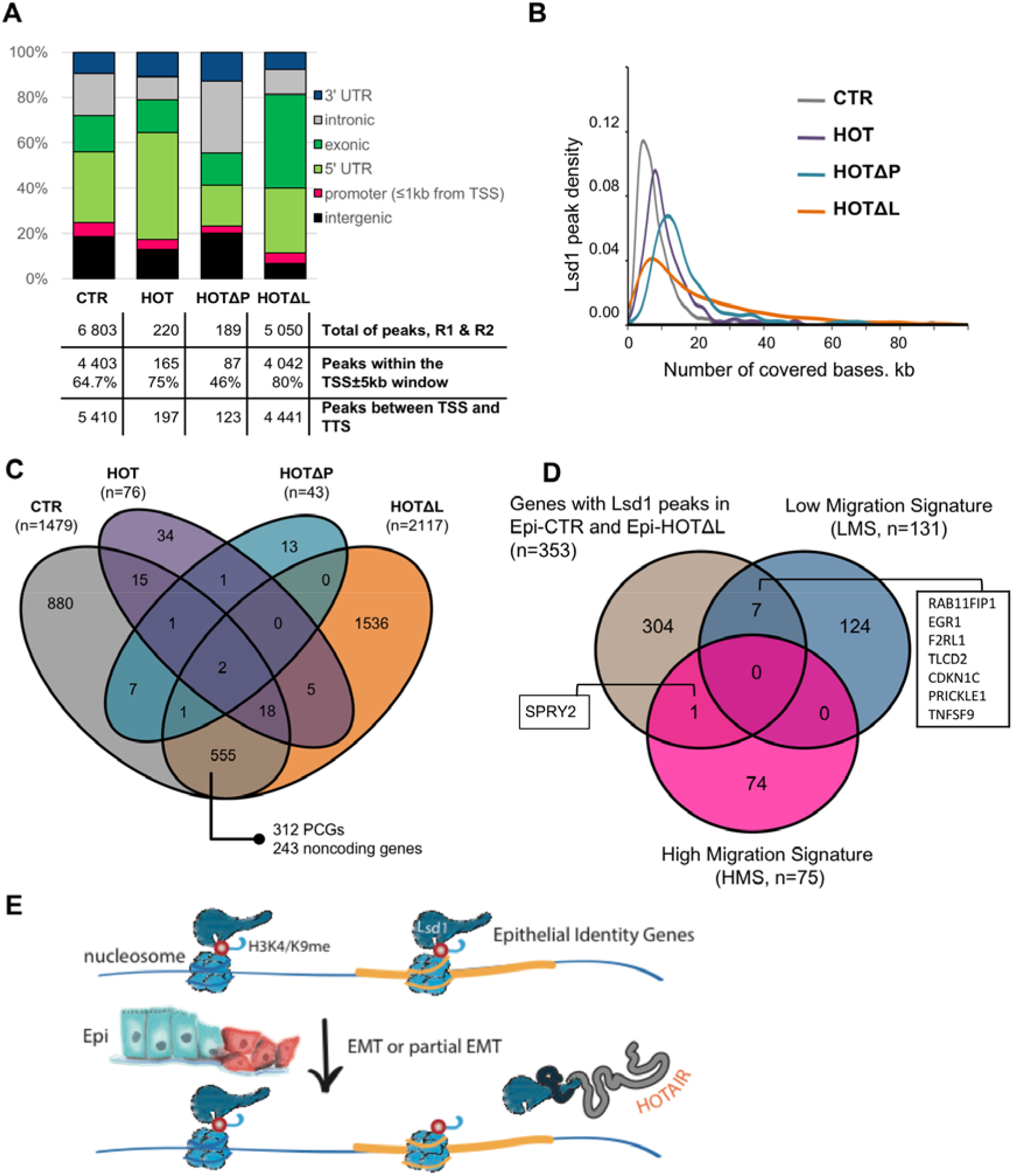
HOTAIR expression promotes Lsd1 dislocation from inherent genomic locations through its 3’-Lsd1-interacting domain: **(A)** Lsd1 peaks identified by ChIP-seq of Lsd1 and SICER peak calling protocol and their distribution across distinct genomic features; **(B)** Density plot of the number of bases covered by Lsd1 peaks; **(C)** Venn diagram representing intersections of genes possessing Lsd1 peaks within the 5 kb window upstream their TSS and within the gene body in Epi-CTR and cells expressing HOT, HOTΔP and HOTΔL; **(D)** Venn diagram presenting the intersection of genes with TSS-associated Lsd1 peaks found in low migrating Epi-CTR and Epi-HOTΔL cells with LMS and HMS sets; **(E)** Model illustrating HOTAIR-mediated disruption of Lsd1 function as a guardian of epithelial identity.

Finally, we determined the number of covered bases per peak for every condition and revealed that all three cell lines expressing HOTAIR showed broader Lsd1 peaks than Epi-CTR (p-value < 10^−11^, Wilcoxon test) (Figure S5D), nevertheless according to the density plot, the two low migrating cell lines, Epi-CTR and Epi-HOTΔL, presented more sharp peaks with the mode values of 3.6 and 6 kb, respectively, comparing to Epi-HOT and Epi-HOTΔP with the mode values of 8.8 and 13.4 kb, respectively (Figure 4B).

Although Lsd1 is not a strictly a promoter associated factor and can be found in distal, enhancer regions, gene bodies, but also cover large chromosomal regions, we decided to explore the Lsd1 landscape on a gene-based approach. For this, we assigned peaks, unique to each cell line and common for two replicates, within the 5kb window around the TSS and within the TSS-TTS window to a corresponding gene and searched for specific Lsd1 patterns associated with low and high migration phenotypes but also with transcriptomic high- and low-migration signatures (HMS and LMS) retrieved from the RNA-seq differential expression analysis (Figure 4C and 4D, Figure S5). Firstly, intersection of the Lsd1-associated genes between conditions revealed high cell-line specificity of Lsd1 loci with only one gene (RNU2-38P, snRNA gene) common to Epi-HOT and Epi-HOTΔP and 555 genes shared by Epi-CTR and Epi-HOTΔL (Figure 4C). We considered the common Lsd1 associated genes of Epi-CTR and Epi-HOTΔL datasets as genomic locations independent of Lsd1/HOTAIR interactions and specific to the low migration phenotype. Among them, 312 represented PCGs and 243 were noncoding genes. Gene enrichment analysis revealed Jak-STAT signaling pathway as the significantly enriched KEGG pathway and several biological processes tightly linked to EMT, such as positive regulation of cell motion and TGF-beta signaling, cytokine-mediated signaling pathways among PCGs presenting Lsd1 peaks 5kb upstream of their TSS and within the gene body (Figure S5E).

Secondly, intersection of Lsd1-associated protein-coding genes (n=353) with LMS (n=131) and HMS (n=75) transcriptomic sets revealed the presence of Lsd1 for 7 up-regulated genes, including the *PRICKLE1*, and for only 1 down-regulated genes, the antagonist of fibroblast growth factor pathways SPRY2, in Epi-CTR and Epi-HOTΔL cells (Figure 4D).

In sum, HOTAIR expression in epithelial cells dramatically affects Lsd1 genomic localization and, in particular, results in its dislocation from specific genomic locations. As a consequence, this imbalances transcription and promotes expression of mesenchymal genes endowing partial transition of epithelial cells to a more mesenchymal phenotype (Figure 4E).

## DISCUSSION

The EMT program is proposed as a one route for generation of normal and neoplastic epithelial cells. It enables acquisition of mesenchymal traits promoting migration and invasion, thus, underlying high metastatic potential of tumor tissues. HOTAIR and Lsd1 have been independently studied in a variety of cell-based and clinical settings as factors associated with EMT and cancer metastasis. And if the regulatory function of HOTAIR is unambiguously linked to acquisition of mesenchymal traits as migration and invasion capacities, the role of the Lsd1 histone demethylase is rather context-specific and resumes in positive or negative control of a variety of cell identity programs. Being ubiquitously expressed in both epithelial and mesenchymal cells, it can induce epigenetic changes either locally (enhancers, promoters, gene bodies) or broadly (large chromatin domains, LOCKs) to influence the transcriptional program both way, through repression or activation (Shi et al., 2004), (Whyte et al., 2012), (McDonald et al., 2011), (Li et al., 2016), (Wang et al., 2007). Lsd1 presence at regions with increased gene expression suggests its positive role through the control of H3K9 methylation status of genes or in tethering of transcription factors promoting transcription initiation (Metzger et al., 2005), (Yang et al., 2019), (Zhang et al., 2018), (Zhang et al., 2018). Another possibility is that Lsd1 regulates co-transcriptional splicing through H3K9 demethylation as it has been shown for CD44 and FGFR2 transcripts (Saint-André et al., 2011), (Gonzalez et al., 2015). Remarkably, in the latter case chromatin modifications have been triggered by the FGFR paired antisense lncRNA (asFGFR2) interacting in cis with PRC2 and KDM2a complexes (Gonzalez et al., 2015). Otherwise, we cannot exclude an indirect effect that Lsd1 may exhibit through a control of upstream factors resulting in up-regulation of LMS genes in Epi-CTR and Epi-HOTΔL cell lines. Further experiments are required to discriminate between these hypotheses.

Lsd1 has also been reported to act independently of its demethylase function at chromatin level and elsewhere (Sehrawat et al., 2018), (Lan et al., 2019), (Carnesecchi et al., 2017). All these mechanistic modalities may be affected by lncRNAs, as described for HOTAIR in breast cancer cells.

The present work identifies HOTAIR as an effector of Lsd1 function as a guardian of epithelial identity. We demonstrated that the 3’-extremity of HOTAIR, which interacts with Lsd1, was essential to promote epithelial cells migration whereas the PRC2-interacting domain was dispensable for this function. Paradoxically and in the light of our results, PRC2 and Lsd1-interacting domains contribute together, but also separately to mechanistically distinct HOTAIR functions. In particular, deletion of the Lsd1-interacting domain still allows cells expressing HOTAIR to maintain a gene expression balance in favor of the epithelial cell identity. This is most likely due to Lsd1 operating independently of HOTAIR. In support of this hypothesis, our Lsd1 chromatin profiling experiments revealed considerable changes in Lsd1 genomic distribution induced by HOTAIR variants with the intact 3’-end sequence enabling its association with Lsd1. The striking correlation between the loss of epithelial traits and changes in Lsd1 landscape strongly supports a pivotal role for Lsd1 as a factor preventing cells from sensing or undergoing an EMT. In the context of effective HOTAIR/Lsd1 association, several molecular scenarios could be considered: (i) HOTAIR may modulate Lsd1 catalytic activity or capacity to interact with its protein partners such as transcription factors or chromatin modifying enzymes (ribo-repressor or activator functions); (ii) HOTAIR may promote the assembly of another specific Lsd1 complex and its tethering to peculiar genomic locations for local chromatin modifications (guide and scaffold functions). Although further studies are required to identify epigenetic changes induced by the Lsd1/HOTAIR complex, to determine other Lsd1/HOTAIR partners and to enlarge this observation to other biological systems, our report revealed an unexpected role of HOTAIR as a molecular toggle switch for Lsd1 function, which may contribute to EMT at the very early steps of transformation of a normal epithelial cell to a neoplastic one.

Intriguingly, several alternative splicing and TSS isoforms of HOTAIR are annotated in the human genome, including those lacking the 3’-terminal sequence interacting with Lsd1 and variants missing the PRC2- or both PRC2- and Lsd1-interacting domains (Mercer et al., 2012). Even if the clinical relevance of these isoforms has not yet been established, in light of our results, one can anticipate that tumors expressing 3’-end truncated variants of HOTAIR would have a lower metastatic potential, and hence, better prognosis. It will be worth assessing the expression of HOTAIR isoforms in tumors of different grades and prognosis to support our findings.

### Experimental Procedures

#### Cell lines and culture

Cells were cultured in a humidified 5% CO_2_ atmosphere at 37°C in high-glucose DMEM with GlutaMAX for HEK293T and MEM alpha without nucleosides for HEK cells (Epi, Mes, Epi-HOT, HOTΔP, HOTΔL) both supplemented with 10% fetal bovine serum, essential amino acids and sodium pyruvate. All cell lines were systematically tested and negative for mycoplasma.

#### Plasmids and oligonucleotides

The CTR construct corresponds to pLenti CMV GFP (659-1) (Addgene, #17445). The HOTAIR full-length and truncated cDNA sequence was PCR amplified from the LZRS-HOTAIR plasmid (Gupta et al., 2010). PCR products were cloned into pENTR and further sub-cloned into pLenti CMV Blast DEST (706-1) (Addgene, #17451) (Campeau et al., 2009) using the Gateway system (Thermo Fisher Scientific). Oligonucleotides sequences for PCR are available in Table S2.

#### Cell transfection and transduction

HEK293T cells at 50-70% of confluence at T25 flasks were co-transfected with 1.3 μg of psPAX2 (Addgene, #12260), 0.8 μg of pVSVG (Addgene, #36399) and 0.8 μg of the lentiviral plasmid bearing cDNA of GFP (CTR), full-length (HOT) or truncated HOTAIR (HOTΔP and HOTΔL) and 5 μL of Lipofectamine® 2000 Transfection Reagent according to manufacturer’s protocol (Thermo Fisher Scientific). Virus supernatant was recovered, filtered 48 h post-transfection and added to Epi cells at 50-70% of confluence. After 24 h post-transduction cells were sub-cultured every two days at a 1:4 ratio in 10 μg/ml of blasticidin supplemented MEM alpha medium for one week and then in MEMalpha medium for additional two weeks prior to any experiment.

#### Wound healing assay (WHA)

Cell suspensions containing 2 00 000 cells were seeded to each well of the 6-well plate and left growing to 95-100% of confluence for 24-36 h. After scratching each wound was imaged twice at time 0 and 24 h post-scratch with a Zeiss inverted microscope with a total of 48 high-field units (HFU) per cell line. The invaded area was calculated as a ratio of Migrated Cell Surface Area to Total Surface Area with a web-based image automated analysis software Tscratch (Gebäck et al., 2009). Results are presented as a mean ± SEM for at least 48 HFUs. P-values were calculated using the Student’s t-test.

#### Subcellular fractionation

This protocol was adapted from (Gagnon et al., 2014). All steps were performed on ice with ice-cold buffers supplemented by 20 U/μl SUPERase-IN (Thermo Fisher Scientific) for RNA extraction and 0.1 mM AEBSF (SIGMA) for protein extraction. RNA extraction from all fractions was performed by Trizol (Thermo Fisher Scientific) at 65°C for 5 min with regular vortexing and then purified with miRNeasy kit (Qiagen) according to the manufacturer’s instructions. For protein extraction, chromatin faction was sonicated for 15 min (30s-on/30s-off, “high power”) on the Bioruptor Plus (Diagenode). Protein content was quantified with the Thermo Fisher Scientific™ BCA Protein Assay Kit (Thermo Fisher Scientific).

#### Protein extraction and Western blot

Cells at 80% of confluence were lysed in RIPA buffer (Thermo Scientific) supplemented with 0.1 mM AEBSF (SIGMA). Total of 5 and 10 μg of the whole protein extract were separated on a NuPAGE Novex 4-12% Bis-Tris Protein Gel and transferred using Thermo Fisher Scientific iBlot Dry Blotting System (Thermo Fisher Scientific). After blocking membranes were immunoblotted with the Epithelial-Mesenchymal Transition Western Blot Cocktail (1:2500 dilution, Abcam, ab157392), anti-GAPDH (Millipore, mab374), EMT Sampler kit (Cell signalling, 9782), anti-Lsd1 (Diagenode, C15410067) overnight at 4 °C, than washed and incubated with a corresponding HRP-conjugated secondary antibody (1:5000 dilution) for 1 hour at room temperature and visualized using the SuperSignal West Femto, Dura or Pico Substrate (Thermo Fisher Scientific). Quantification of protein amounts was performed relative to GAPDH using the ImageJ software.

#### Immunofluorescence

Suspension of 20000 cells was seeded on a 4-well microscope slide and let growing for 24 h. Following PBS wash cells were fixed in 3% Formaldehyde-2% Sucrose for 15 minutes, permeabilized and blocked with 10% goat serum (Thermo Fisher Scientific). For F-actin staining, fixed and permeabilized cells were incubated with 0.5 μg/ml DMSO solution of phalloidin-tetramethylrhodamine isothiocyanate (TRITC) (Sigma) for 40 min at room temperature. For ZO-1, cells were incubated with the primary antibody (1:100 dilution, Cell Signaling Technology, 8193) for 40 minutes at 37°C, washed in PBS and incubated with Alexa Fluor 594 Goat anti-Rabbit secondary antibody (1:400, Thermo Fisher Scientific, A11037) for 30 minutes at 37°C. Slides were washed and mounted using a ½ dilution of Vectashield Mounting Medium with DAPI (Vector Laboratories).

#### Fluorescent microscopy and image analysis

For each cell line, three images per well were acquired as stacks composed of twenty slices acquired with an interval of 0.5 μm, each generated from 3 ApoTome images over the height of the cells with a 40X objective on a Zeiss AxioVision microscope equipped with an ApoTome module (PiCT-BDD platform, Institut Curie). All images were captured with identical exposure time for DAPI and Alexa Fluor 594 using AxioVision software and then analyzed using the Fiji software (Schindelin et al., 2012). Stacks were processed with the « Z-project » tool (« Sum Slices » setting) and an identical minimum threshold-value was set for all images. Signal was then quantified using the « Measure » tool and normalized by the number of cells in the image. Integrated density per cell was again normalized on overall image background signal to reduce hybridization bias between slides and presented as a mean ± SEM for at least 11 power field units.

#### RNA extraction and reverse transcription

Total RNA extraction was performed directly from cell cultures with miRNeasy kit according to the manufacture’ instructions (Qiagen). Only RNAs with the RNA Integrity Number (RIN) above 6 were used for further experiments. Reverse Transcription (RT) was performed on 500 ng of RNA with either random and oligo(dT) primers mix (iScript™ cDNA Synthesis Kit) or specific oligonucleotides (SuperScript II Reverse Transcriptase, Thermo Fisher Scientific) (Table S2). Reactions without reverse transcriptase were included as a negative control for DNA contamination.

#### RNA-seq library preparation

1 μg of RNA was depleted for ribosomal RNA with the RiboMinus™ Eukaryote Kit for RNA-seq (Thermo Fisher Scientific) and converted into cDNA library using a TruSeq Stranded Total Library Preparation kit (Illumina). cDNA libraries were normalized using an Illumina Duplex-specific Nuclease (DSN) protocol prior to a paired-end sequencing on HiSeq™ 2500 (Illumina). At least 20x coverage per sample was considered as minimum of unique sequences for further data analysis. Raw RNA-seq data is available at Gene Expression Omnibus (GEO) under accession number GSE106517.

#### RNA-seq data analysis

Reads were mapped allowing 3 mismatches using TopHat 2.0.4 (Trapnell et al., 2009) and the human genome version hg19. Uniquely mapped reads were assembled using the BedTools suite (Quinlan and Hall, 2010) and merged in segments if mapped in the same strand to the Gencode V15 annotation to extract protein-coding genes and annotated noncoding genes including lncRNA, antisense, sense_intronic, sense-overlapped and pseudogenes.

Differential expression analysis was performed using the DESeq R package (Anders and Huber, 2010). Only genes with an expression fold change (FC) above 2 and adjusted p-value below 0.05 were retained as significantly differentially expressed. Gene ontology (GO) biological process, KEGGS pathways and cellular compartment terms enrichment analysis was performed using DAVID 6.8 and REVIGO (Huang et al., 2009), (Huang et al., 2008), (Supek et al., 2011).

#### Chromatin immunoprecipitation (ChIP)

Total of 10^6^ cells from 80% confluent culture was used per IP, crosslinked in 1% formaldehyde for 10 minutes at 37 °C, scraped and lysed in Cell Lysis Buffer (5 mM PIPES pH 8, 85 mM KCl, 0.5% NP-40). The lysate was passed through a 20G long needle 10 times and nuclei were recovered by centrifugation at 400g for 15 min at 4°C. The nuclei pellet was resuspended in Nuclear Lysis Buffer (50 mM Tris-HCl pH 8, 10 mM EDTA pH 8, 1% SDS) and sonicated in Bioruptor Plus (Diagenode) for 30 minutes at 4 °C. Following centrifugation fragmented chromatin was diluted to a final concentration of 0.1% SDS, pre-cleared with 20 μl of BSA blocked Protein A beads (Thermo Fisher Scientific) at 4°C for 2 hrs. Total of 1/10 volume of the pre-cleared chromatin was taken as the Input-DNA sample and the rest was incubated with a 1 μg of the primary antibody (Diagenode) for 12 h at 4°C in a rotating wheel. The chromatin was immunoprecipitated by addition of 50 μl of BSA blocked beads for 2 more hours at 4°C, washed twice with FA150 (20 mM Tris-HCl pH 8, 150 mM NaCl, 2 mM EDTA pH 8, 1% Triton X-100, 0.1% SDS), twice with FA500 (the same as FA150, except 500 mM NaCl), and once with TE (10 mM Tris-HCl ph8, 1 mM EDTA). Chromatin was eluted in Elution Buffer (10 mM Tris-HCl pH8, 1% SDS) at 65°C for 1 hour. Input and IP chromatin were further reverse cross-linked and DNA was purified by QIAquick PCR Purification Kit (Qiagen) with 30 μl of final elution volume. DNA concentration of IP- and Input-DNA was measured by Qubit with dsDNA HS Assay Kit (Thermo Fisher Scientific).

#### ChIP-seq library preparation and sequencing

ChIP-seq libraries were prepared with the TruSeq ChIP-Sample Prep Kit-Set A and PCR Box (Illumina) from 5 ng of IP- and Input-DNA. Library was amplified by 10 PCR cycles following the manufacture’s procedures. Its concentration and quality was assessed with Qubit with dsDNA HS Assay Kit (Thermo Fisher Scientific) and High Sensitivity DNA Analysis Chip Kit (Agilent Technologies). Libraries were sequenced on a HiSeq 1500 device in a fast single-end mode and 50 bp read length. A range of 10-25 million of reads passing Illumina quality filter was obtained for each sample. Raw ChIP-seq data is available at GEO under accession number GSE106262.

#### ChIP-seq data analysis

The reads were first aligned to human genome version hg19 using the bowtie algorithm (Langmead and Salzberg, 2012). Only those tags that mapped to unique genomic locations with at most one mismatch were retained for further analysis. The peaks were called from uniquely mapped reads by the SICER v1.1 algorithm with redundancy threshold 1, window size 200, effective genome fraction 0.74, gap size from 400 to 1600 for the first test and then equal to 1000 for the final analysis (Zang et al., 2009). Peaks were then annotated with gene information, including the distance to TSS and locations in gene function elements (intron, exon, UTR, etc.) from genome mapping information of RefSeq transcripts using HOMER v4.1 software (Heinz et al., 2010). Distance between the middle of each LSD1 peak and the transcript start site (TSS) and the transcription termination site (TTS) coordinates of known genes was calculated using Gencode v26 annotation.

#### Gene features analysis

Gene enrichment analysis was done using DAVID (Huang et al., 2008), (Huang et al., 2009) and enrichment score was calculated as −log10(p-value).

#### Quantitative PCR

cDNA was diluted 10-50 times and quantified by real-time PCR with LightCycler® 480 SYBR Green I Master (SIGMA). The RPL11 gene was used as a reference to quantify relative abundance of each cDNA. Error bars represent standard deviations in relative quantity of cDNA prepared from at least three independent experiments.

---

**Figure S1.**
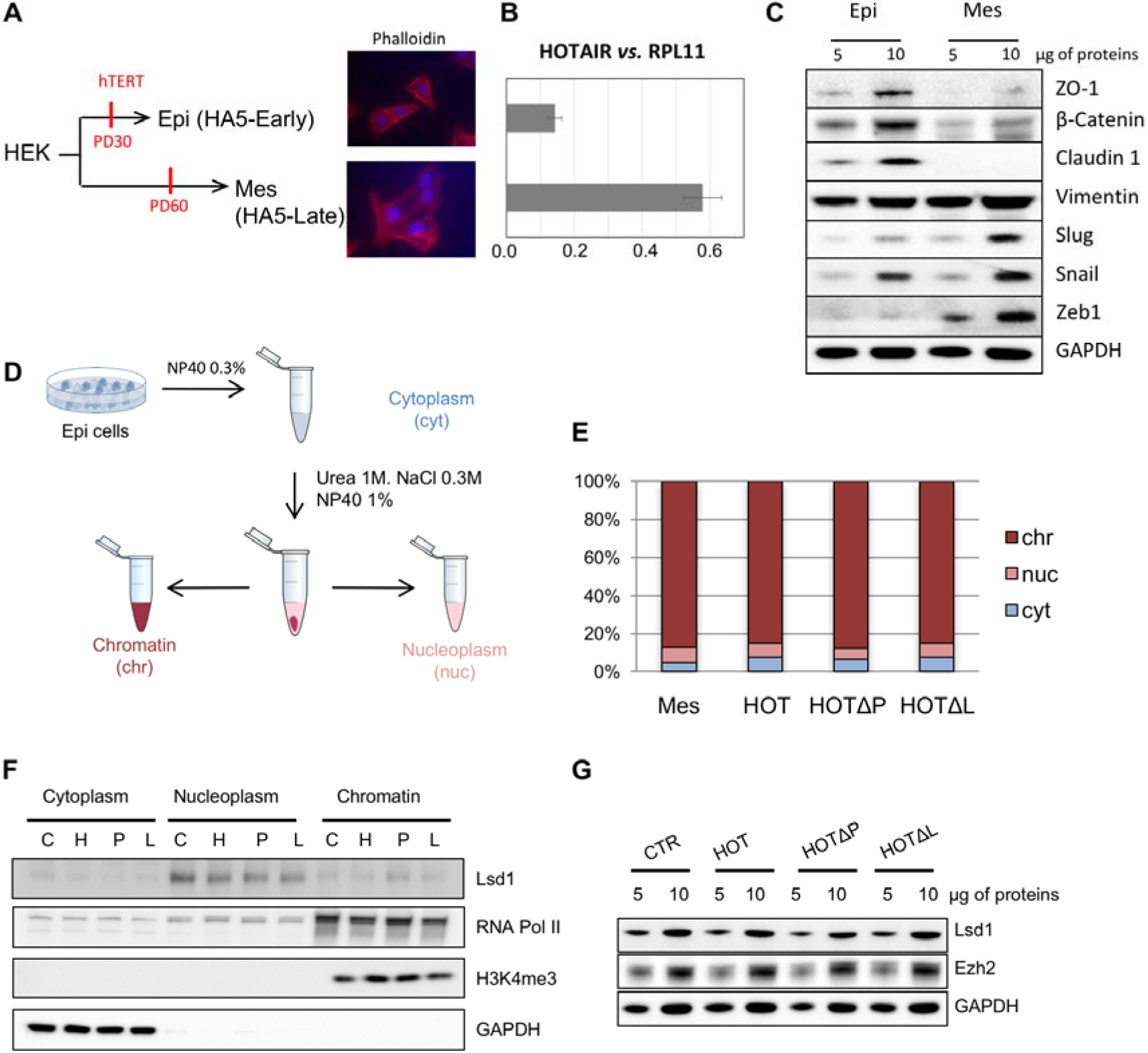
In *vitro* EMT system used to study HOTAIR function: **(A)** Epi and Mes cell lines corresponding to HA-Early5 and HA5-Late, respectively (Castro-Vega et al., 2013), stained for F-actin fibers by Phalloidin-TRITC (x40); **(B)** Quantification of HOTAIR expression in Epi and Mes cells by random-primed RT-qPCR; **(C)** Expression levels of EMT markers in whole protein extracts of Epi and Mes cells assessed by Western blot; **(D)** Protocol of subcellular fractionation into cytoplasm, nucleoplasm and chromatin fractions; **(E)** Distribution of full-length and truncated variants of HOTAIR between cytoplasm, nucleoplasm and chromatin fractions in Mes and Epi cells assessed by RT-qPCR relative to GAPDH mature mRNA; **(F)** Subcellular distribution and levels of Lsd1, RNA Pol II, H3K4me3 and GAPDH in cytoplasm, nucleoplasm and chromatin fractions and **(G)** Lsd1, EZH2 and GAPDH levels in whole protein extracts assessed by Western blot in Epi cells expressing CTR (C), HOT (H), HOTΔP (P) and HOTΔL (L).

**Figure S2.**
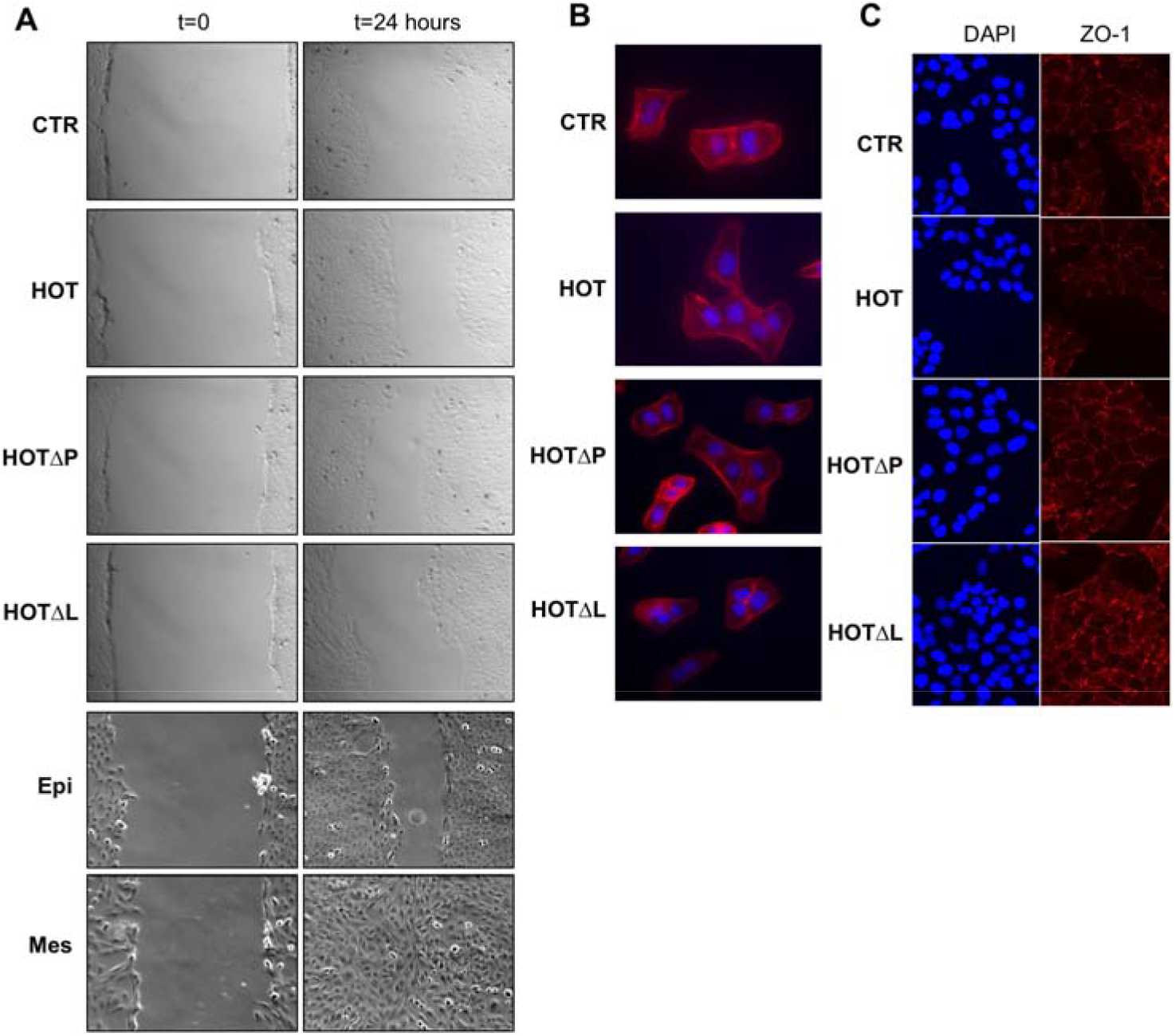
EMT characteristics of Mes and Epi cells used in this study: **(A)** Assessment of migration capacities by WHA: **r**epresentative images at zero and 24 hours post-scratch; **(B)** Phalloidin-TRITC staining of F-actin fibers (x40); **(C)** ZO-1 subcellular localization assessed by Immunostaining and fluorescence microscopy: representative images of ZO-1 (red) and DNA/nucleus (DAPI, bleu) generated from three ApoTome stacks in Epi cells expressing none (CTR), full-length (HOT) and truncated variants (HOTΔP and HOTΔL) of HOTAIR.

**Figure S3.**
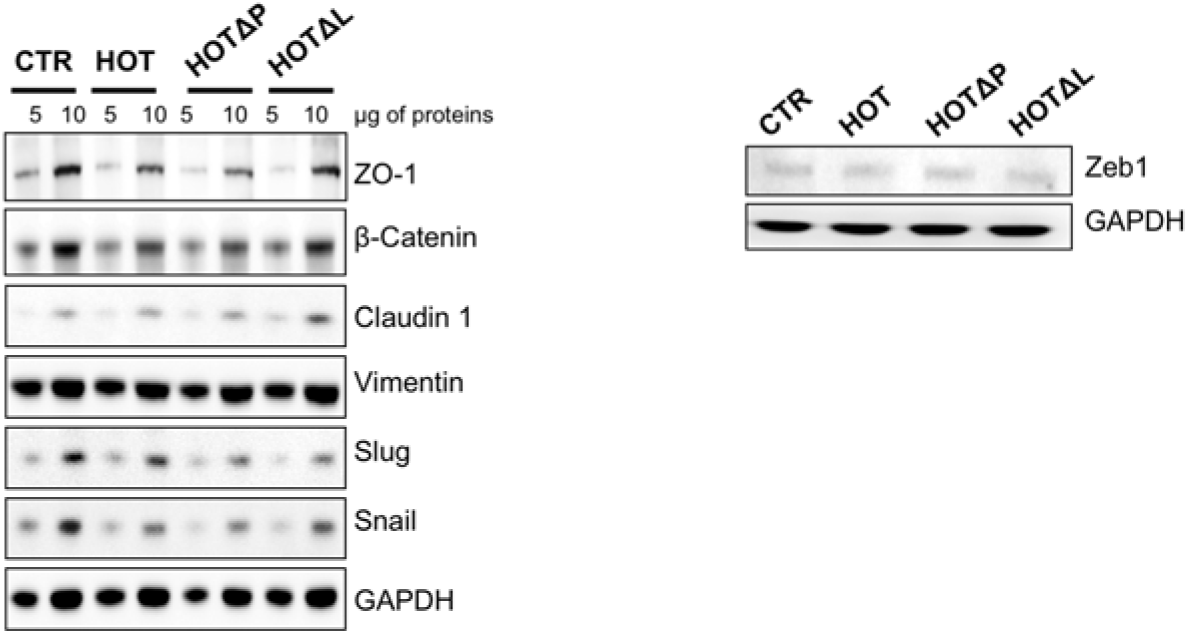
Expression levels of EMT markers in whole protein extracts of Epi-CTR, HOT, HOTΔP or HOTΔL cell lines assessed by Western blot.

**Figure S4.**
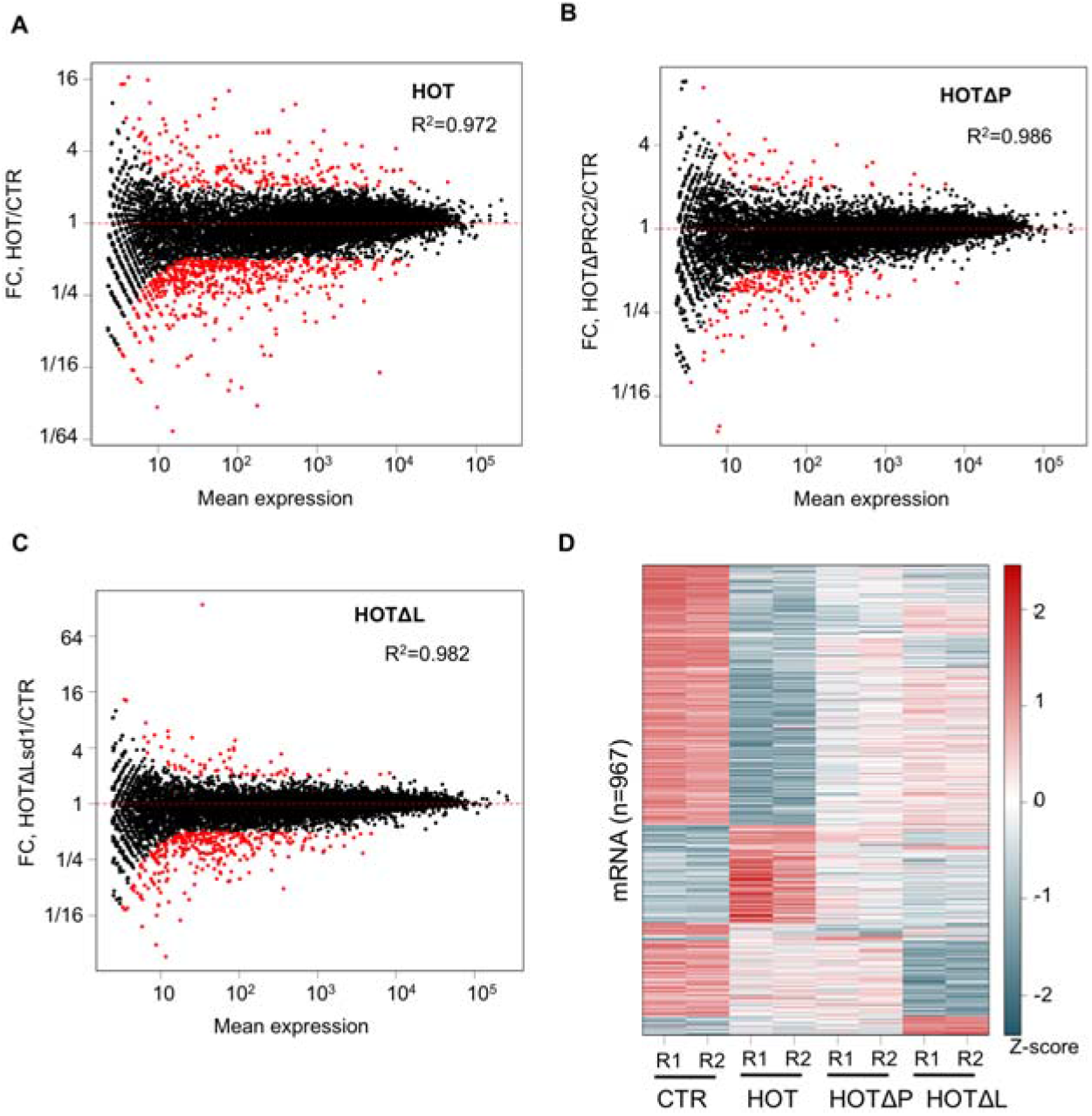
Expression of full-length and Lsd1-interacting domain deleted variant of HOTAIR induces the most drastic changes in PCG transcriptome: **(A-C)** MA-plot of protein-coding genes expression in Epi-HOT, HOTΔP and HOTΔL cells comparing to Epi-CTR; Black dots represent all counted PCG, red dots only those with the fold-change (FC) above 2 and adjacent p-value below 0.05; **(D)** Heatmap of DE-PCGs defined by RNA-seq and DESeq analysis.

**Figure S5.**
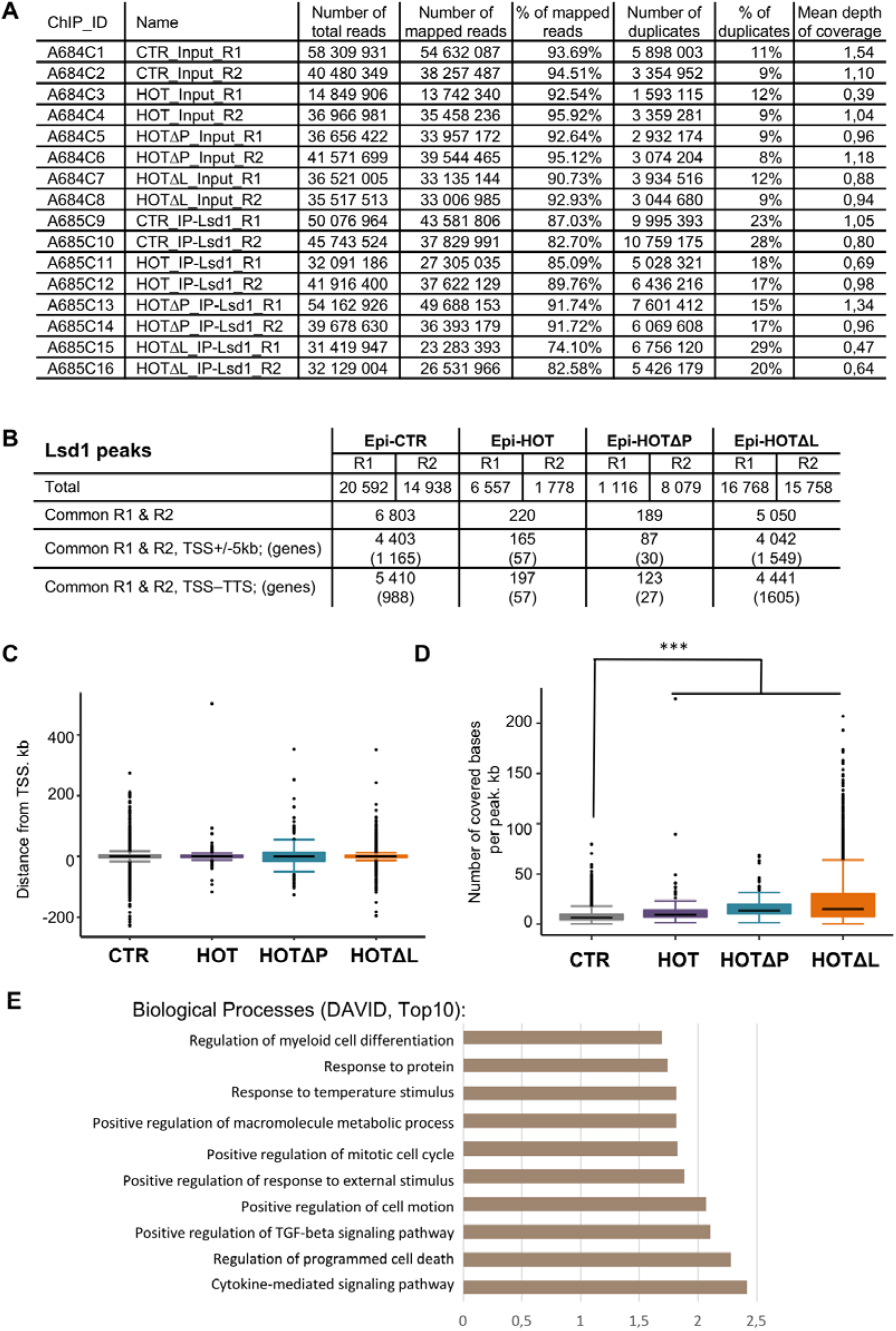
Lsd1 peak features: **(A)** ChIP-seq metrics of Input- and IP-DNA sequencing; **(B)** SICER identified Lsd1 peaks in Epi-CTR (n=6803), HOT (n=220), HOTΔP (n=189) and HOTΔL (n=5050) cells, per replicate and common between two replicates; **(C)** Box plot of Lsd1 mead peak distances from genes TSS in Epi-CTR (n=6803), HOT (n=220), HOTΔP (n=189) and HOTΔL (n=5050) cells; **(D)** Box plot of number of bases covered by Lsd1 peaks in Epi-CTR (n=6803), HOT (n=220), HOTΔP (n=189) and HOTΔL (n=5050) cells: *** p-value < 10^−11^, Wilcoxon test; **(E)** TOP10 hits of biological processes identifies by DAVID as significantly enriched within the set of PCGs presenting Lsd1 peaks within the 5kb window upstream their TSS and within the gene body.

**Table S1.**
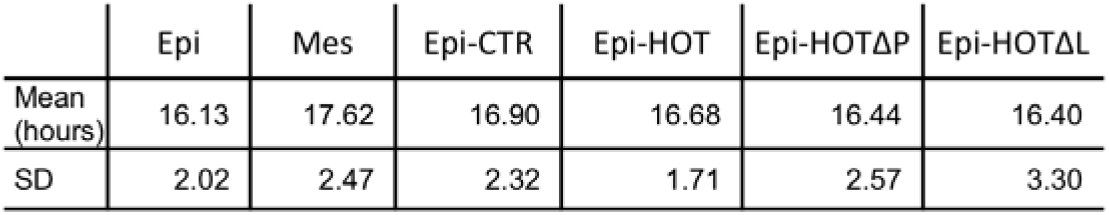
Population doubling (PD) time expressed as a mean with a standard deviation (SD) calculated for exponentially growing cells according to a formula PD=(t_F_-t_I_)*ln2/ln(N_F_/N_I_); t stands for time; F and I stand for Final and Initial, and N stands for the number of cells.

## Author Contributions

Project administration and funding acquisition, A.M., M.P.; conceptualization, supervision and writing, M.P.; draft revisions, D.F., J.J.; investigation, M.P., C.B., J.J., D.F.; formal analysis, M.G., Z.S., M.D.; expertise, A.L-V..

**The authors declare no conflict of interest.**

## Acknowledgments

We deeply thank Marc Descrimes, Elena Battistella and Sandra Carignon for important technical contributions, Karina Jouravleva and Luis Castro-Vega for fruitful discussions and expertise sharing, and all members of Morillon’s lab for reading and commenting the manuscript. The HOTAIR cDNA containing LZRS plasmid was a kind gift of H.Y. Chang.

