## Supplementary material for "HOTAIR promotes an epithelial-to-mesenchymal transition through relocation of the histone demethylase Lsd1": Table S2

**Table S2.** Oligonucleotides used in this study for cloning and RT-qPCR.

| gene | Application | sense | ID | sequence, 5'-3' |
|---|---|---|---|---|
| HOTAIR | qPCR | frw | AMOh16 | GGTAGAAAAAGCAACCACGAAGC |
| HOTAIR | RT and qPCR | rev | AMOh17 | GTGAGTGCCCGTCTTGCCCT |
| HOTAIR | Gateway clonning | frw | AMOh353 | GGGGACAAGTTTGTACAAAAAAGCAGGCTGACTCGCCTGTGCTC TGGAGCTTGATCCGA |
| HOTAIR | Gateway clonning | rev | AMOh354 | GGGGACCACTTTGTACAAGAAAGCTGGGTTTTTTTTTTTGAAAAT GCATCCAGATATTA |
| HOTdP | Gateway clonning | frw | AMOh475 | GGGGACAAGTTTGTACAAAAAAGCAGGCTTCTTTATTTTTTTAAG GCC |
| HOTdP | Gateway clonning | rev | AMOh476 | GGGGACCACTTTGTACAAGAAAGCTGGGTTTATATTCACCACAT GTAAAA |
| HOTdL | Gateway clonning | frw | AMOh477 | GGGGACAAGTTTGTACAAAAAAGCAGGCTCCAGTTCTCAGGCGA G |
| HOTdL | Gateway clonning | rev | AMOh478 | GGGGACCACTTTGTACAAGAAAGCTGGGTTGGTTTCACTTTTAAA AATTT |
| GFP | Gateway clonning | frw | AMOh367 | GGGGACAAGTTTGTACAAAAAAGCAGGCTAAAGGAGAAGAACTT TTCAC |
| GFP | Gateway clonning | rev | AMOh368 | GGGGACCACTTTGTACAAGAAAGCTGGGTTTTGTATAGTTCATCC ATGC |
| RPL11 | qPCR | frw | AMOh172 | AGCAGCCAAGGTGTTGGAG |
| RPL11 | RT and qPCR | rev | AMOh173 | TACTCCCGCACCTTTAGACC |
| CD44 | qPCR, exon2 | frw | AMOh60 | TGCCGCTTTGCAGGTGTAT |
| CD44 | RT and qPCR, exon2 | rev | AMOh61 | GGCCTCCGTCCGAGAGA |
| CD44 | qPCR, exon14 | frw | AMOh1597 | TGGAAGATTTGGACAGGACAGG |
| CD44 | RT and qPCR, exon15 | rev | AMOh1598 | AGAAGCTCTGAGAATTACTCTGC□ |
| PRICKLE1 | qPCR | frw | AMOh670 | GACAGTCTCTCCTCTTATCG |
| PRICKLE1 | RT and qPCR | rev | AMOh671 | CTCTGCCTTTCCAAAATTCTTCAC |
| MTUS1 | qPCR | frw | AMOh666 | AGCTTCGGGACACTTACATT |
| MTUS1 | RT and qPCR | rev | AMOh667 | ATAGGCCTTCTTTAGCAATTC |
| EGR1 | qPCR | frw | AMOh697 | CTTCAACCCTCAGGCGGACA |
| EGR1 | RT and qPCR | rev | AMOh698 | GGAAAAGCGGCCAGTATAGGT |
| TGFB2 | qPCR | frw | AMOh616 | CCAAAGGGTACAATGCCAAC |
| TGFB2 | RT and qPCR | rev | AMOh617 | CAGATGCTTCTGGATTTATGGTATT |
| CCND1 | qPCR | frw | AMOh1588 | GTGTGCAGAAGGAGGTCCTGC□ |
| CCND1 | RT and qPCR | rev | AMOh1590 | CCTCCTCGCACTTCTGTTCC□ |
| DNER | qPCR | frw | AMOh620 | ATGCCAGTTCTAACAGCTCTGC |
| DNER | RT and qPCR | rev | AMOh621 | GGAGCACTGTTGGAATCCTGTGG |
